# Anomalous Diffusion Characterization by Fourier Transform FRAP with Patterned Illumination

**DOI:** 10.1101/2020.03.13.990937

**Authors:** A.C. Geiger, C.J. Smith, N. Takanti, D.M. Harmon, M.S. Carlsen, G.J. Simpson

## Abstract

Fourier transform fluorescence recovery after photobleaching (FT-FRAP) with patterned illumination is theorized and demonstrated for quantitatively evaluating normal and anomalous diffusion. Diffusion characterization is routinely performed to assess mobility in cell biology, pharmacology, and food science. Conventional FRAP is noninvasive, has low sample volume requirements, and can rapidly measure diffusion over distances of a few micrometers. However, conventional point-bleach measurements are complicated by signal-to-noise limitations, the need for precise knowledge of the bleach beam profile, potential for bias due to sample heterogeneity, and poor compatibility with multi-photon excitation due to local heating. In FT-FRAP with patterned illumination, the time-dependent fluorescence recovery signal is concentrated to puncta in the spatial Fourier domain through patterned bleaching, with substantial improvements in signal-to-noise, mathematical simplicity, representative sampling, and multiphoton compatibility. A custom nonlinear-optical beam-scanning microscope enabled patterned illumination for photobleaching through two-photon excitation. Measurements in the spatial Fourier domain removed dependence on the bleach profile, suppressing bias from imprecise knowledge of the point spread function. For normal diffusion, the fluorescence recovery produced a simple single-exponential decay in the spatial Fourier domain, in excellent agreement with theoretical predictions. Simultaneous measurement of diffusion at multiple length scales was enabled through analysis of multiple spatial harmonics of the bleaching pattern. Anomalous diffusion was characterized by FT-FRAP through a nonlinear fit to multiple spatial harmonics of the fluorescence recovery. Constraining the fit to describe diffusion over multiple length scales resulted in higher confidence in the recovered fitting parameters. Additionally, phase analysis in FT-FRAP was shown to inform on flow/sample translation.

**Statement of Significance:** Fourier transform fluorescence recovery after photobleaching (FT-FRAP) with patterned illumination greatly improves the accuracy of diffusion assessments and simultaneously accesses information on both normal and anomalous diffusion in a single experiment.

## 1. Introduction

Fluorescence recovery after photobleaching (FRAP) is a well-established and widely accessible method for probing diffusion.^1-2^ In FRAP, a region of a fluorescently labeled sample is permanently photobleached using a short, high-intensity burst of light. After the bleach, mobile fluorescent molecules diffuse into the region and mobile bleached molecules diffuse out of the region. This combined mobility results in a time-dependent recovery of fluorescence intensity in the bleached region. Diffusion information can be obtained by fitting the fluorescence recovery to a mathematical model.

The first FRAP experiment was performed by Peters et. al. in 1974 to measure the mobility of membrane proteins in red blood cell ghosts.^3^ More recently, FRAP has been used to probe epidermal growth factor receptor clustering in Chinese hamster ovary cell membranes,^4^ intercellular communication via septal junctions in multicellular cyanobacteria,^5^ and the dynamics of intermediate filament-like protein in the hyphae of Streptomyces venezuelae.^6^ FRAP has also been applied broadly in the pharmaceutical community to understand molecular transport in hydrogels^7-10^ and in extracellular matrices^11-12^ in an effort to improve drug delivery outcomes. Recent efforts have also been made to utilize FRAP as a pre-screening assay for *in meso* crystallization of membrane proteins in lipid cubic phase.^13-16^

Despite the advantages of FRAP, quantitative diffusion analysis is typically complicated by the requirement for precise knowledge of the bleaching profile.^17-18^ To support rapid diffusion measurements, FRAP is generally optimized for fast recovery by using a small bleach spot. To compensate for low signal from a small bleach spot, high bleach depth is used to increase the signal-to-noise ratio (SNR). However, increasing the bleach depth runs the risk of complicating reproducibility in the spatial bleach profile by introducing nonlinearities from optical saturation and perturbations to diffusion from local heating.^19^ To address this issue, alternative bleach patterns have been explored, most notably disc^20^, line^21^, and fringe pattern^22-24^ illumination. Disc illumination has the advantage of increasing the overall number of molecules bleached, but largely negates the 1/f noise reductions from highly localized bleaching and correspondingly fast recoveries. Line illumination is a reasonable compromise, supporting fast recovery in the direction orthogonal to the bleach line and signal to noise averaging along the length of the line. With the possible exception of disc and fringe illumination, in which the contiguous bleach spot is large relative to the optical point-spread function, the point and line bleach patterns with the greatest reduction in 1/f noise are most prone to artifacts from ambiguities in the bleach point spread function (PSF).

Spatial Fourier analysis (SFA) is one of the more successful strategies used to date for addressing ambiguities in the PSF for point-excitation.^20, 25-27^ In summary, diffusion in FRAP can be modeled as the convolution of the bleach PSF with a time-varying Gaussian function. This convolution produces a function describing the real-space fluorescence recovery that is dependent on both time and the bleach PSF. Moreover, this convolution generally produces a real-space recovery with no simple closed-form analytical solution, with a few notable exceptions for the PSF (e.g., Gaussian). However, in the Fourier transform domain, the convolution corresponds to a simple multiplication, disentangling the time-dependent decay from the bleach profile. The decay curves for each spatial frequency in SFA can be used individually or collectively for recovering the diffusion coefficient. In this manner, the detailed functional form for the initial PSF become less critical in the analysis, as the fluorescence recovery is only dependent on time and not on the initial bleach PSF for a single spatial frequency.^28^ For point-excitation, SFA suffers by distributing the signal power from sharp features in the real-space image out over many low-amplitude frequencies in the SFA image, but the intrinsic SNR can be recovered through simultaneous, collective analysis at multiple spatial frequencies. When one or a small number of frequencies are used, this distribution of power can result in a reduction in SNR, the cost of which represents a trade-off for the benefits in reducing ambiguities related to the PSF.

Furthermore, point-bleach FRAP lacks sensitivity for characterizing anomalous diffusion. Diffusion is categorized as anomalous when it deviates from normal Brownian diffusion. Whereas the mean squared displacement (MSD) in normal diffusion evolves with a linear dependence on time, the MSD in anomalous diffusion exhibits a nonlinear dependence on time and/or space, resulting in a time-varying/distance-dependent diffusion coefficient.^29-30^ Anomalous diffusion has been observed in a variety of systems, such as the cell and polymeric networks.^31-34^ Anomalous diffusion in point-bleach FRAP can be identified through a nonlinear fit of the fluorescence recovery to an anomalous diffusion model.^35^ However, the relatively subtle differences in the point-bleach recovery curves between normal and anomalous diffusion can complicate identification and quantification of deviations from normal diffusion. Further complications in accurately characterizing anomalous diffusion with point-bleach FRAP arise from the requirement for precise knowledge of the bleaching PSF. In addition, significant covariance between fitting parameters can result in relatively large uncertainties in the recovered coefficients (e.g., the diffusion coefficient and an anomalous exponent).

Finally, point excitation poses particularly problematic practical challenges from local heating effects in multi-photon excited FRAP measurements.^36^ Because of the general inefficiency of multi-photon excitation, a large flux of light is typically introduced, only a small fraction of which contributes to excitation and fluorescence. Weak but nonzero absorption of the incident light and Stokes Raman transitions leading to local heat deposition both compete with multi-photon excitation. When the excitation beam is fixed at a single location, local temperatures can quickly escalate until the rate of heat dissipation matches the rate of deposition. Depending on the steady-state temperature differential, this transient temperature gradient can potentially bias subsequent diffusion measurements based on isothermal assumptions.

In this work, comb patterns for illumination during photobleaching were demonstrated to support high SNR measurements of normal and anomalous diffusion in Fourier transform analysis of the fluorescence recovery with multi-photon excitation. In brief, bleach patterns were selected to concentrate signal to puncta in the spatial Fourier transform domain, rather than a point in the real-space image, as shown in **Figure 1**. Patterned illumination using rapid line-scanning distributed the power from the bleach over much larger regions in the field of view, removing many of the potential nonlinearities and biases associated with highly localized excitation while enabling multi-photon excitation with negligible artifacts from local heating. Probing diffusion at multiple length scales by interrogating multiple harmonics in the spatial frequency domain was shown to increase confidence in recovered fitting parameters in analysis of anomalous diffusion. The theoretical foundation for Fourier-transform fluorescence recovery after photobleaching (FT-FRAP) with patterned illumination is evaluated in proof-of-concept studies of model systems for characterizing normal and anomalous diffusion.

**Figure 1.**
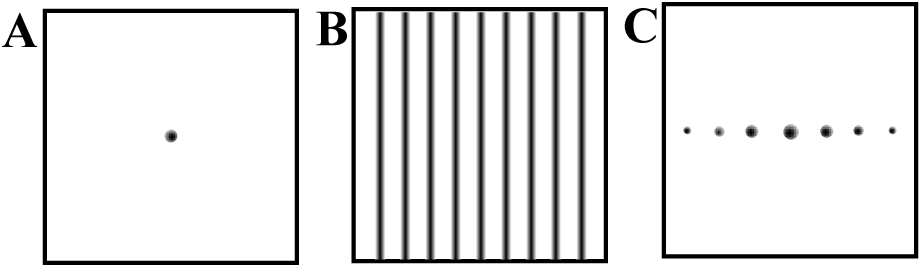
Schematic depicting A) conventional FRAP illumination, B) patterned illumination with comb excitation, and C) Spatial FT of patterned illumination in B. Conventional FRAP produces a sharp point in real space. FT-FRAP with patterned illumination produces sharp puncta in the spatial Fourier domain.

## 2. Theoretical Foundation

### 2.1 Fourier Analysis of the Diffusion Equation

Prior to discussion of FT-FRAP, it is useful to review conventional FRAP as a comparator within the FT framework. Fick’s Law of Diffusion is given by the following general expression. 

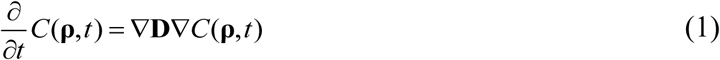

In which **C**(**ρ**,*t*) is the concentration of the analyte of interest as a function of position **ρ** and time *t*, and **D** is the three-dimensional diffusion tensor.

The differential equations within the diffusion equation are arguably simpler to evaluate by first performing the spatial Fourier transform to generate 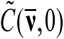, in which 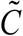 is the spatial Fourier transform of *C* and 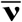 is the 3D spatial wavevector. Upon Fourier transformation of the diffusion equation given in **Equation (1)**, each derivative transforms as multiplication by the “diagonal” function 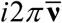. 

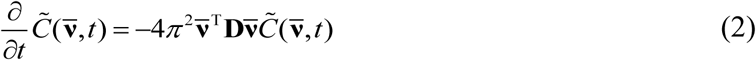

Evaluation of 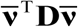 considering diffusion just within the (*x,y*) plane yields the nonzero scalar products 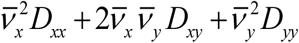. For a choice of (*x,y*) coordinates defined along the principal moments of the diffusion tensor **D** (including the case of constant diffusion in all (*x,y*) directions), *D*_*xy*_ = 0. In this case, the diffusion equation can be independently evaluated in each of the (*x,y*) directions. The expression for the *x*-direction is given below, with an analogous expression present for the *y*-direction. 

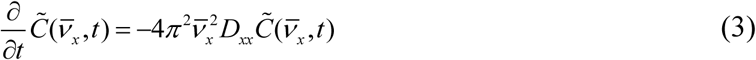

Since only 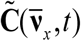 in **Equation (3)** depends on time, the expression in **Equation (3)** is of the form *f’*(*t*) = *kf*(*t*), for which one general solution, subject to the constraint of a decay, is given by an exponential function of the form *f*(*t*) = *f*(0)*e*^*kt*^, with *k* < 0. 

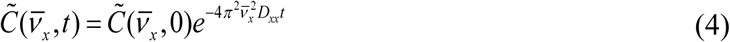

This solution to the differential equation along the *x* direction is a Gaussian function in 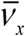. Multiplication by a Gaussian in the spatial Fourier domain corresponds to convolution with a Gaussian in real space. 

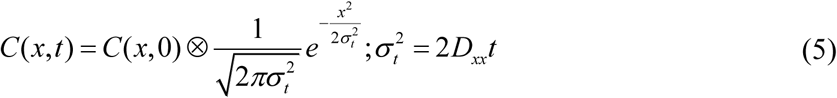

The standard deviation of the spatial Gaussian distribution *σ*_*t*_ increases with the square root of time, corresponding to convolution with an ever-broadening Gaussian as diffusion proceeds.

### 2.2 Conventional FRAP

In the limit of a thin sample, diffusion in the *z*-direction can be neglected, with diffusion expressed with respect to both *x* and *y*. Under these conditions in conventional FRAP (depicted in **Figure 1A**), the bleach pattern for a symmetric Gaussian illumination pattern on the back of an objective is also Gaussian within the field of view. 

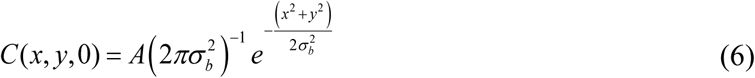

In isotropic media, diffusion is identical in both the *x* and *y* coordinates, such that the diffusion tensor can be replaced by a single scalar diffusion coefficient *D*. In this limit, the convolution of two 2D Gaussian functions (one from the initial bleach and one from diffusion) has the convenient property of producing yet another Gaussian function, the width of which evolves in time. 

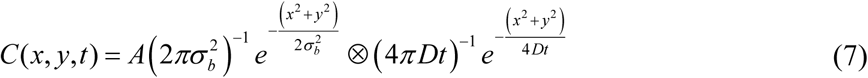

Evaluation of the convolution results in the following expression for the time-dependence. 

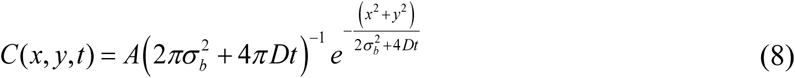

If the system is anisotropic, selection of the principal spatial coordinates that diagonalize the diffusion matrix allows the diffusion equation to incorporate differences in diffusivity along different spatial dimensions. 

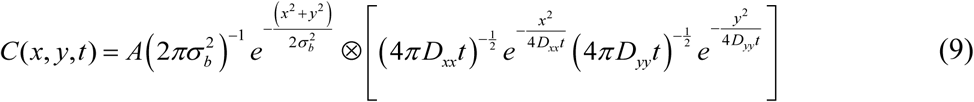

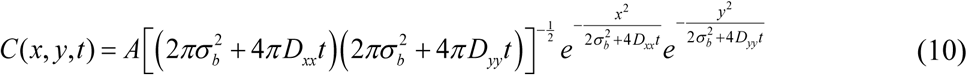

In the more general case of a non-Gaussian function describing the bleach pattern *C*(*x,y*,0), the situation is significantly more complex. In general, no simple analytical forms are expected for the convolution of a Gaussian with non-Gaussian functions, requiring numerical methods for approximations. Unfortunately, non-Gaussian bleach patterns are commonplace. Even when Gaussian patterns are intended, bleaching can often approach saturating conditions when the peak bleach depth approaches unity, resulting in “top-hat” initial bleach peak shapes. In such cases, the shape of the recovered region can be complicated to integrate analytically into the diffusion analysis for recovery of the diffusion coefficient.

### 2.3 Comb Bleach FT-FRAP

In FT-FRAP, the initial bleach pattern is selected to produce sharp puncta in the spatial frequency domain, rather than in real-space. One such pattern is a comb, or a periodic series of lines. For mathematical purposes, we will define the comb pattern to proceed along the *x*-axis in the laboratory frame with constant illumination along the *y-*axis, producing a series of bleached stripes (depicted in **Figure 1B**). For a bleaching PSF along the *x*-axis of *PSF*(*x*), the initial bleach pattern *C*(*x*,0) is constant in the *y*-axis and given below, in which 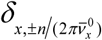 is a delta function at the positions 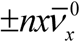 and *n* ∈ {0,1, 2…}, *C*_*0*_ is the initial concentration of the analyte of interest, and *A* is the bleach depth. 

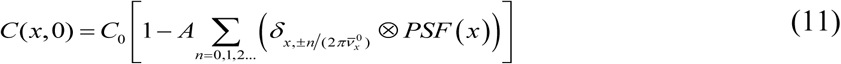

By taking the spatial Fourier transform of this equation, the convolution of the bleach *PSF(x)* with the comb pattern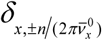 is replaced by a multiplication operation, simplifying the analysis. The initial (*t* = 0) spatial Fourier transform of *C*(*x*,0) along the *x*-direction is then given by the following expression, in which 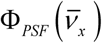 is the spatial Fourier transform of *PSF*(*x*). 

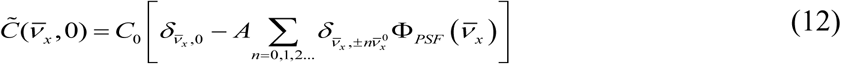

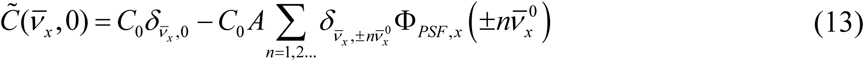

In brief, the initial bleach corresponds to a series of puncta in the spatial Fourier domain positioned at 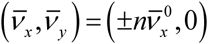, each of which is scaled in initial amplitude by the spatial Fourier transform of *PSF*(*x*), with additional amplitude at the origin 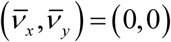 from the overall average fluorescence intensity.

The time-dependent behavior of each punctum in the FT can be evaluated by an approach analogous to that illustrated in **Equation (4)** for the *n*th harmonic. 

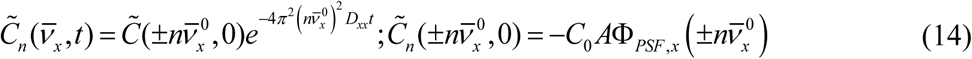

For a given impulse in the FT image, a single-exponential decay is expected; irrespective of the functional form for *PSF*(*x*), the time-constant is given by 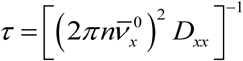. This single-exponential recovery is in stark contrast to conventional FRAP analysis based on measurements performed in real-space, for which the time scale for recovery depends sensitively on precise foreknowledge of *PSF*(*x*) for recovery of the diffusion coefficient. The time-constant of the fluorescence recovery in FT-FRAP is independent of *PSF(x)*, which allows for simplified mathematical recovery of *D*_*xx*_ while circumventing error associated with imprecise estimates of *PSF(x)*.

Through this analysis, comb bleaching has the additional practical advantage of simultaneously enabling diffusion analysis over multiple length scales. For example, by nature of the quadratic dependence on spatial frequency, the fourth harmonic will recover 16-fold faster than the first harmonic for normal diffusion. This disparity enables analyses of both rapidly and slowly diffusing species without the need for changing instrument settings.

### 2.4 Anomalous Diffusion

The capability of FT-FRAP to simultaneously measure diffusion over multiple length scales enables quantitative analysis of anomalous diffusion. While normal diffusion is characterized by fluorescence recovery with a quadratic dependence on spatial frequency, anomalous diffusion will produce harmonics with fluorescence recovery that deviates from this dependence. The ability to simultaneously interrogate diffusion over several discrete, well-defined distances by FT-FRAP with comb illumination provides a convenient route for quantifying anomalous diffusion, if present.

Numerous mathematical models for anomalous diffusion can be found for trends anticipated under a diverse suite of conditions.^30, 37-41^ A model based on continuous-time random walk and fractional diffusion is considered in this work because of its general applicability to systems with both time-varying and distance-dependent diffusion.^29^ Deviation from normal Brownian diffusion can arise from various sources, two of which we will consider in the anomalous diffusion model used in this work. First, anomalous diffusion can arise when there is a distribution in the “wait times” between each step. If diffusion is described as a series of random steps, normal diffusion has a constant time between each step, resulting in an MSD with a linear time dependence. When the distribution of wait times broadens due to binding or association, the time dependence of the MSD deviates from linear and scales with time to the power *α*. 

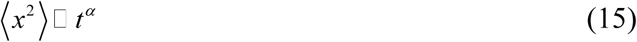

Normal diffusion corresponds to *α =* 1, subdiffusion corresponds to 0 < *α* < 1, and superdiffusion corresponds to *α* > 1. The fluorescence recovery of a sample in this class is described by a one-parameter Mittag-Leffler function E_*α*_.

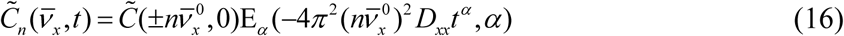

The Mittag-Leffler function is a fractional generalization of an exponential function. The one-parameter Mittag-Leffler, *E*_*α*_ (*z,α*) converges to an exponential *E* (*z*,1) = *e*^*z*^ for *α =* 1, and 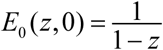 for *α* = 0.

A second source of anomalous diffusion is a distribution in step length. Normal diffusion exhibits a constant step length leading to a quadratic dependence of the diffusion coefficient on spatial frequency. However, in cases where there is a broad distribution of step lengths, Lévy flight behavior is observed and the anomalous diffusion coefficient will adopt a dependence on *σ*^*µ*^ instead of *σ*^*2*^ for normal diffusion, where *σ* is the standard deviation of the step-length distribution and *τ* is the characteristic waiting time. 

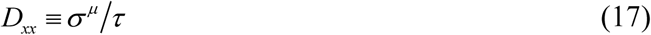

Lévy-flight diffusion produces a stretched exponential decay in the spatial frequency domain, where the spatial frequency is raised to the power *µ* rather than 2 for normal diffusion. 

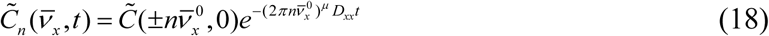

A system can exhibit both subdiffusive and Lévy-flight behavior when there is are broad distributions in both wait times and in step lengths. Modification of the Mittag-Leffler function shown in **Equation (16)** by replacing the quadratic dependence on the spatial frequency term with the exponent *µ* and by scaling the exponent on *t* by *2/µ* results in an equation that can describe deviation from normal Brownian diffusion in both time and space. 

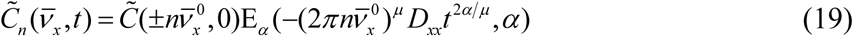

FT-FRAP is capable of sensitively characterizing anomalous diffusion because it can measure diffusion on multiple length scales. A fit to the equations describing anomalous diffusion involves parameters that are likely to have high covariance if the fit is performed on only one recovery curve. By performing a global fit with multiple recovery curves at multiple length scales, the fit can be constrained to recover more accurate values for the parameters describing anomalous diffusion.

### 2.5 Signal Power

In comparison to analysis in real space, the Fourier domain analysis with patterned illumination provides a substantial advantage in terms of the available power of the detected signal. Power is conserved upon Fourier transformation, allowing direct comparisons across both representations. For a Gaussian bleach spot with a width parameter *σ*, and peak bleach depth *A*_*p*_, the power is generated by integration over the 2D Gaussian. 

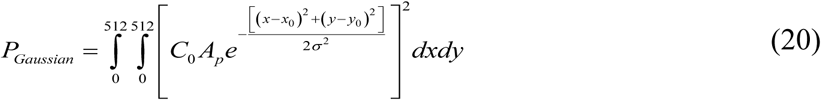

In a typical experiment, the Gaussian width is much smaller than the field of view in order to reduce measurement times by minimizing the diffusion length. In the limit that *σ* <<512 pixels (assuming a field of view of 512×512 pixels), the discrete limits of integration can be safely evaluated as ±∞. The integrals can be further simplified by substituting *x*′ = *x*-*x*_0_ and noting that *dx*=*dx*′ (with analogous substitutions for *y*). 

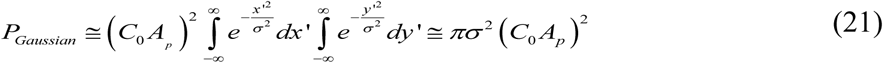

The power in the impulse in the FT image produced by comb illumination can be similarly calculated by evaluating the same power through integration of the real-space bleach pattern. For a bleach depth *A*_*p*_, the power in a comb with *N* teeth is given by the following expression. 

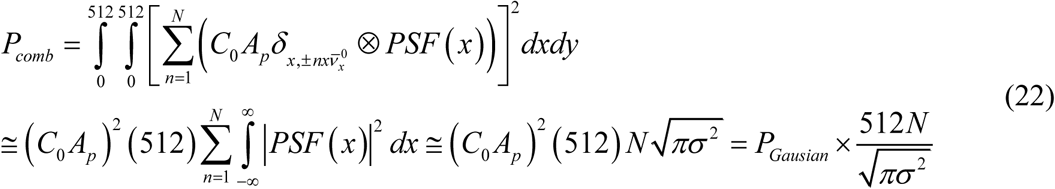

The power advantage for comb excitation with a beam PSF with a characteristic width of 2 pixels relative to point excitation is ∼2,300-fold greater than that in the real-space spot with an identical bleach depth and 32 “teeth” in the comb. This potential for signal-to-noise enhancement is particularly noteworthy since the peak bleach amplitude *A*_*p*_ is bounded to be less than unity and is typically much less than 0.5 to reduce nonlinear effects from saturation and local heating.

### 2.6 Phasor Analysis of Flow in FT-FRAP

The preceding description is based on the assumption of even functions for the comb bleach pattern relative to the origin of the image (typically, the center). Even assuming the initial bleach pattern is symmetric about the origin, directional flow within the sample could result in displacements along the flow direction (assumed to be x for simplicity) over time. Displacements in real space correspond to shifts in phase in the Fourier domain, such that phase analysis in the FT domain has the potential to inform on flow. Considering comb excitation, a shift of Δ*x* in the initial bleach pattern will produce shifts in the *δ*-functions associated with the comb. 

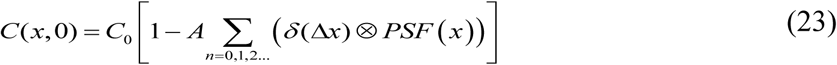

In which 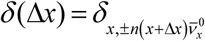. The influence of displacement is easily integrated into the FT analysis using the shift theorem, in which displacement by an offset from the origin of Δ*x* is accounted for in the spatial FT through multiplication by 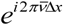 for a given value of *n.*^42^ 

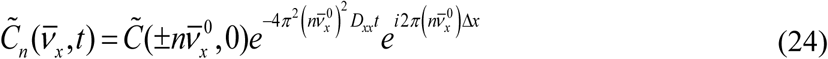

At *t* = 0, the initial phase angle of the *n*th reflection is related to the argument *φ*_*n*_ of 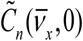, which is simply 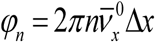. In the absence of time-averaged flow in the *x*-direction, the argument of the *n*th reflection will be preserved throughout the experiment. If flow is nonzero, then Δ*x* is a function of time. Assuming a constant flow rate of *q*_*x*_ = Δ*x*/*t*, then Δ*x* can be replaced by *q*_*x*_*t* + Δ*x*_0_ in **Equation (24)**, in which Δ*x*_0_ is the phase shift at *t* = 0. This substitution results in an argument for the *n*th peak given by the following equation. 

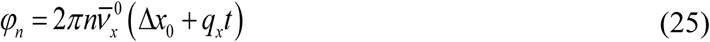

Notably, the rate of change in the phase shift from flow is proportional to *n*, such that the higher harmonics corresponding to higher spatial frequencies are likely to be more sensitive to flow than the lower harmonics. This trend mirrors analogous sensitivities to time in the fluorescence recovery from diffusion in the preceding section, in which the higher harmonics report on fast diffusion times measured over short distances.

## 3. Experimental Methods

### 3.1 Two-photon excited fluorescence (TPEF) microscopy

The experimental apparatus for two-photon excitation in FT-FRAP is depicted in **Figure 2**. A tunable 80 MHz, Ti:sapphire, femtosecond laser (Mai Tai) purchased from Spectra-Physics (Santa Clara, CA) was used for the excitation source. The fundamental beam was raster-scanned across the sample using a 8.8 kHz resonant scanning mirror purchased from Electro-optical Products Corporation (Ridgewood, NY) for the fast-scan axis and a galvanometer mirror purchased from Cambridge-Tech (Bedford, MA) for the slow-scan axis, both controlled by custom timing electronics built in-house.^43^ A 10X, 0.3 NA objective purchased from Nikon (Melville, NY) was used to focus the beam onto the sample, and the TPEF signal was collected in the epi direction through the same objective used for delivery of the excitation beam. The laser was tuned to 800 nm with a power of ∼50 mW at the sample during imaging and ∼500 mW at the sample during bleaching. A long-pass dichroic mirror (650DCXR) purchased from Chroma (Bellows Falls, VT) and a band-pass filter (FGS900) purchased from Thorlabs (Newton, NJ) were used to isolate the TPEF signal before it was detected by a photomultiplier tube (PMT) (H7422P-40 MOD) purchased from Hamamatsu (Hamamatsu City, Shizuoka, Japan). Responses of the PMT were digitized synchronously with the laser pulses by using a digital oscilloscope card (ATS9350) purchased from AlazarTech (Pointe-Claire, Québec, Canada) and mapped onto 512 × 512 images via custom software written in-house using MATLAB purchased from MathWorks (Natick, MA).^43^ The TPEF videos were recorded at ∼4 frames per second.

**Figure 2.**
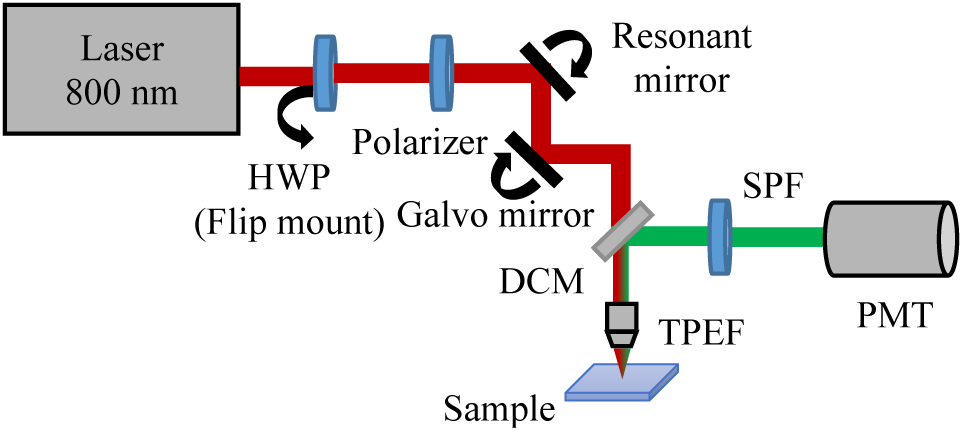
Instrument schematic of home-built nonlinear-optical beam-scanning microscope used for multi-photon FT-FRAP. HWP on flip mount is used to modulate from low power (∼50 mW) to high power (∼500 mW) for the bleach. Beam scanning is performed with galvanometer (slow axis) and resonant (fast axis) mirrors. DCM = dichroic mirror, HWP = half-wave plate, PMT = photomultiplier tube, SPF = short-pass filter, TPEF = two-photon excited fluorescence.

### 3.2 Comb bleach FT-FRAP

A simple change to the scan pattern of the galvanometer (slow axis) mirror was used to generate a comb bleach pattern at the sample. Following an initial low-power period for baseline TPEF microscopy of the full field of view, patterned bleaching was performed simply by changing the number of steps in the ramp function driving the galvanometer mirror from 512 (used for normal imaging) to an integer fraction of 512 corresponding to the fundamental spatial frequency (e.g. 8, 16, 32 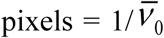). The dwell time per step was also increased proportionally such that the repetition rate of the slow axis mirror was independent of the number of lines in the comb bleach pattern. A flip mount with a half-wave plate (depicted in **Fig. 2**) was synchronized to switch the excitation source from low power to high power concurrently with the reduction in ramp steps. This protocol resulted in a comb bleach pattern as seen in **Figure 3A**. After ∼2 seconds at high power the flip mount was removed, reducing the laser power, and the number of steps for the slow axis mirror was changed back to 512 to facilitate normal imaging at low power to track the fluorescence recovery of the sample.

**Figure 3.**
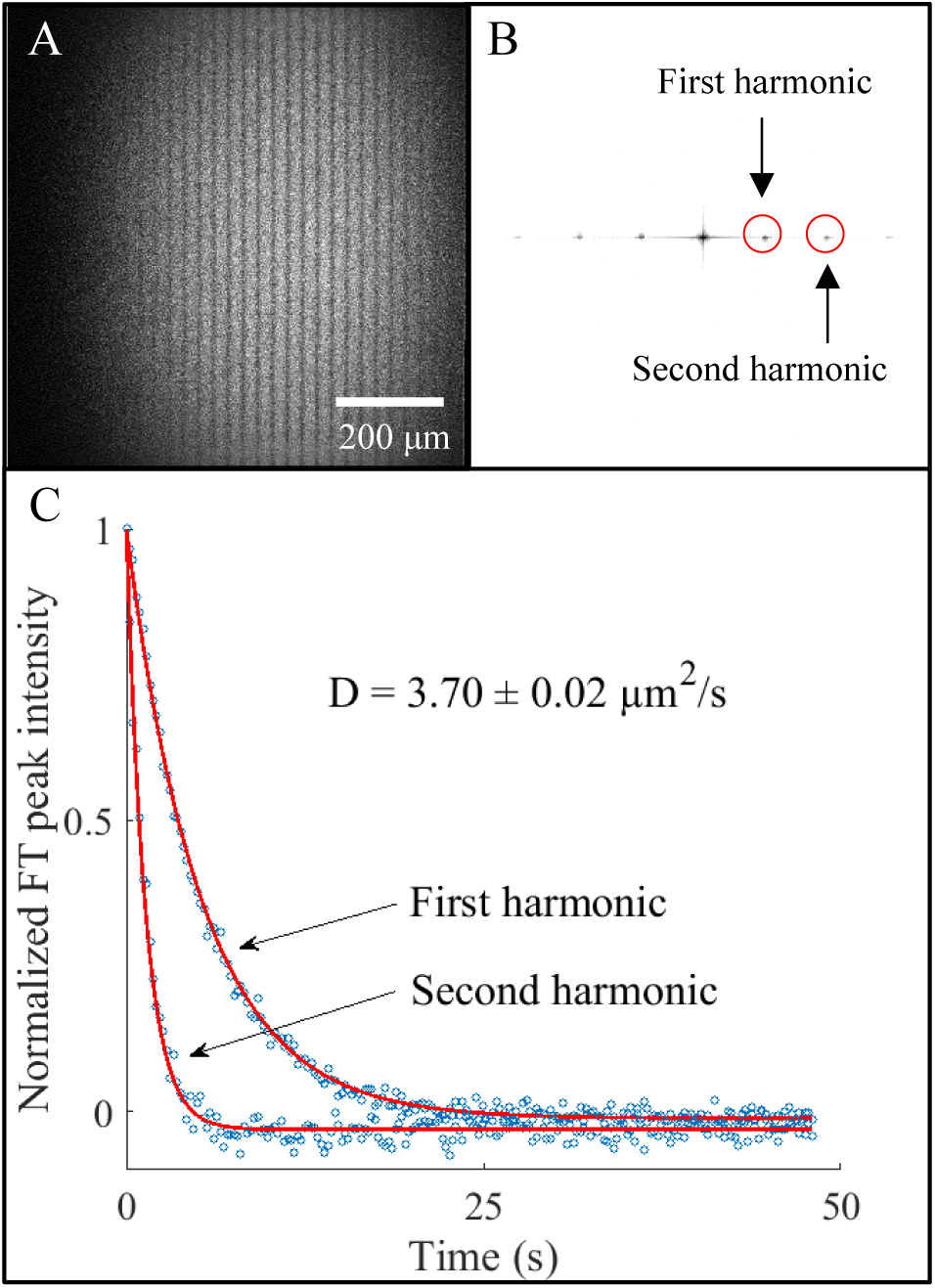
Multiphoton excited FT-FRAP with comb bleach of FITC-polydextran (2 MDa) dissolved in 50/50 glycerol/water. A) Image of the solution immediately after a 32-line comb bleach. B) 2D-FT of (A) with the circled peaks corresponding to the first and second spatial frequency harmonics of the 32-line comb bleach. C) Fluorescence recovery of the first and second harmonic peaks with best-fit curves recovering a diffusion coefficient, *D* = 3.70 ± 0.02 μm^2^/s. The reported uncertainty is the standard deviation of the fit.

### 3.3 Data analysis

Analysis of the FT-FRAP data was performed using custom software written in-house using MATLAB. A 2-dimensional FT was taken of each image. FT-FRAP curves were recovered by integrating over peaks in the FT magnitude. A fit was performed to recover the diffusion parameters using **Equation (14)** for normal diffusion and **Equation (19)** for anomalous diffusion. A MATLAB function written by Roberto Garrappa was used for evaluating the Mittag-Leffler function in **Equation (19)**.^44^ Uncertainties in the fits were calculated based on the second derivative of *χ*^2^-space in the vicinity of the minimum. Phasor analysis of flow was performed by taking the argument of the complex-valued 2D-FT peaks and relating the phase back to flow velocity through **Equation (25)**.

### 3.4 Sample preparation

Solutions of 2 mg/mL fluorescein isothiocyanate (FITC) polydextran (2 MDa) purchased from MilliporeSigma (Burlington, MA) were used to evaluate the FT-FRAP approach. These fluorescently labeled molecules were solubilized in either 50/50 glyercol/water or in an aqueous solution of 22 mg/mL hyaluronic acid (15 MDa) purchased from Lifecore Biomedical (Chaska, MN). Solutions were mixed thoroughly prior to FRAP analysis.

## 4. Results and Discussion

### 4.1 FT-FRAP characterization of normal diffusion

The capabilities of FT-FRAP were evaluated using polydextran, a model system for diffusion measurements. As shown in **Figure 3A**, a comb pattern was employed for photobleaching a solution of 2 mg/mL FITC-polydextran (2 MDa) in 50/50 glycerol/water. Following photobleaching, spatial Fourier transformation produced sharp puncta with symmetric amplitudes about the origin peak as shown in **Figure 3B**. As diffusion progressed, the FT peak intensities exhibited simple exponential decays, as shown in **Figure 3C**. These observations are in excellent agreement with theoretical predictions in **Equation (4)**. Consistent with prior arguments on signal power, the FT-FRAP analysis clearly provides high signal-to-noise bleaching curves by combining the analysis over the entire field of view.

Comb patterns for bleaching enabled simultaneous analysis over multiple length scales through the *n*th spatial harmonics 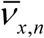, as shown in **Figure 3**. Theoretical predictions in **Equation (14)** suggest an exponential decay of each impulse with a decay constant given by 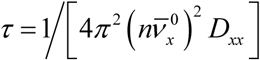. Higher harmonics are expected to exhibit faster decays with a quadratic dependence on *n*. The decays of the first and second harmonic peaks were fit to **Equation (14)**, recovering a diffusion coefficient of 3.70 ± 0.02 µm^2^/s.

### 4.2 FT-FRAP characterization of flow

The impact of flow on the phase of the recovered Fourier transform peaks was evaluated in the results shown in **Figure 4**, in which unidirectional fluid motion was simulated by sample translation with an automated stage during the fluorescence recovery. The sample under investigation was an aqueous solution of 2 mg/mL FITC-polydextran (2 MDa) in 50/50 glycerol water. Recovery of the diffusion coefficient in FT-FRAP only requires analysis of the magnitude of the Fourier peaks. However, the real and imaginary components of the Fourier peaks contain information about the spatial phase of the bleach pattern. **Figure 4A** shows an oscillatory decay of the real and imaginary components of first and second harmonic peaks. The phase of the Fourier peak can be calculated using the argument of the complex number describing the peak at each time point. **Figure 4B** shows the phase shift of a sample that was not translated and the phase shift of the first and second harmonic peaks of a sample that was translated. Consistent with the expression in **Equation (25)**, the phase angle changed linearly in time for the system undergoing directional translation in the *x*-direction of the bleach comb, with a proportionality constant of 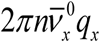 Consequently, the *n*th harmonic is predicted to exhibit an *n*-fold increase in the rate of phase angle change over time relative to the fundamental peak. From the results summarized in **Figure 4B**, precisely this trend was observed in the measurements, in excellent agreement with the theoretical predictions. The measured flow rate for the translated sample (3.49 ± 0.02 μm/s) matched closely with the translation rate of the sample stage (3.33 μm/s) after correcting for bulk flow from convection measured from the experiment in the absence of translation.

**Figure 4.**
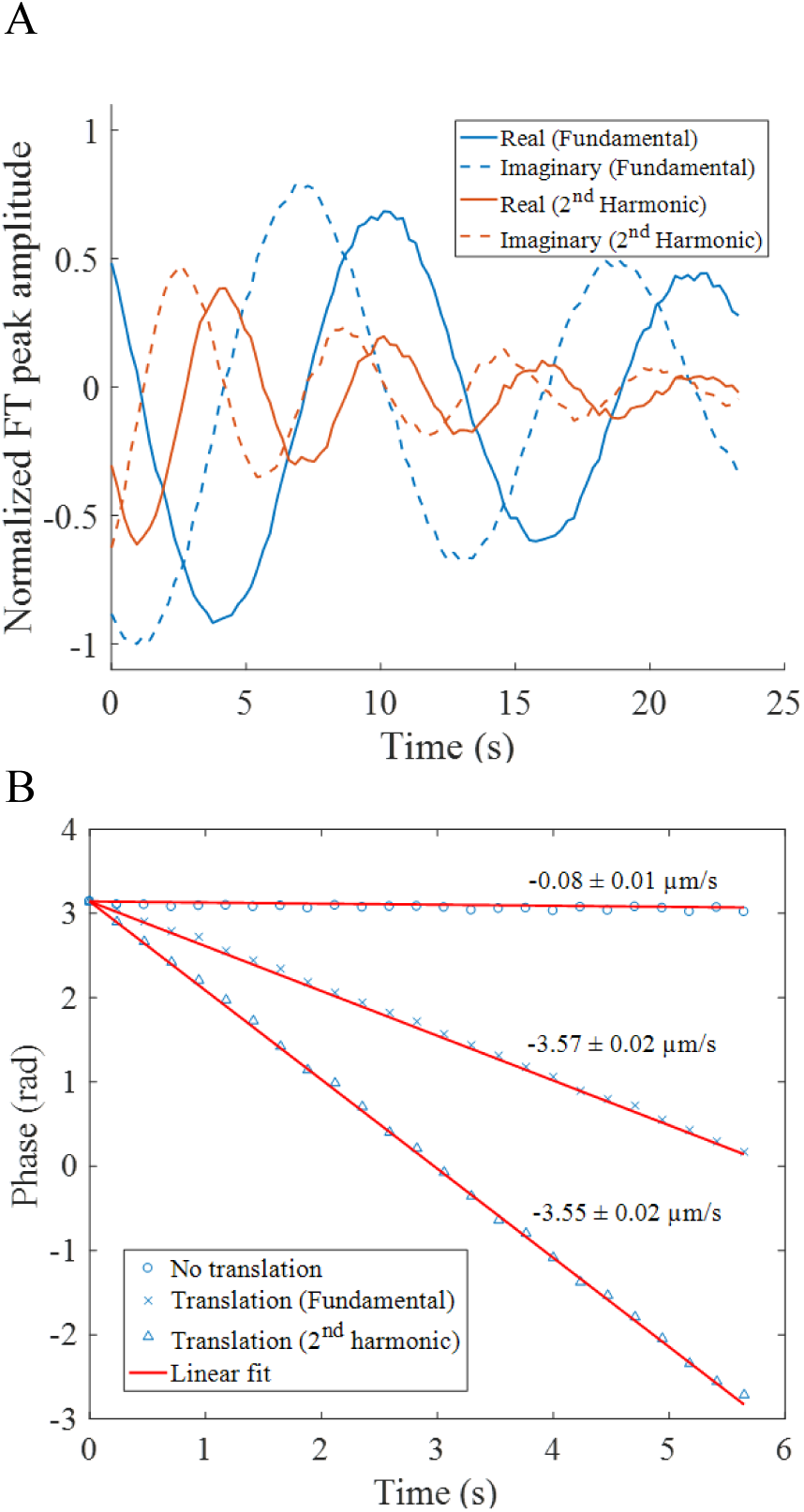
Bulk flow measured by FT-FRAP of FITC-polydextran (2 MDa) in 50/50 glycerol/water. A) Real and imaginary amplitudes of the 2D-FT fundamental and 2^nd^ harmonic peaks upon sample translation during diffusion. B) Phase calculated from real and imaginary amplitudes of 2D-FT peaks. The non-translated sample has minimal bulk flow. The velocities calculated from the fundamental and 2^nd^ harmonic peaks of the translated sample are within one standard deviation of each other and close to the translation rate of the sample stage (3.33 μm/s) after correcting for bulk flow from convection. The reported uncertainties are the standard deviation of the fit.

### 4.3 Patterned versus point bleach profiles

It is worthwhile to compare the FT-FRAP approach with patterned illumination demonstrated herein with previous studies employing spatial Fourier analysis (SFA) of FRAP measurements. In those prior studies, Fourier analysis was performed to aid in interpretation of recoveries using conventional bleach illumination of localized points. SFA provided similar computational benefits in the mathematical simplicity arising in the Fourier domain. However, FT-FRAP has a major signal-to-noise advantage over conventional point-illumination. In the numerator, FT-FRAP supports major increases in the signal power (>2000-fold with comb illumination) by distributing the bleach amplitude over the entire field of view, whereas conventional point-illumination saturates (bleach depth approaching unity) at a much lower integrated signal power. Furthermore, patterned illumination enables shifting of the signal to a quiet spatial frequency for noise suppression. By analogy with 1/*f* noise in electronics, analysis of natural images suggests a power spectrum obeying a 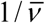 dependence.^45^ For optical detection in the shot-noise limit, the variance in signal is proportional to the mean. Since visible photons are often detectable with signal-to-noise approaching the shot-noise limit in instrumentation optimized for FRAP, it stands to reason that the noise in the Fourier domain will also scale with the signal power in an image with natural contrast. Consequently, the low frequency noise power spectrum is also expected to scale with 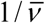, in direct analogy with 1/*f* noise in electronics. As in electronics, shifting of the signal to a frequency regime with lower noise through modulation can provide a substantial noise reduction.

The illumination patterns investigated in this work were specifically designed for spatial Fourier transform analysis and as such differ significantly from a host of previous FRAP studies using patterned illumination. Previous work investigated the use of arbitrary photobleach patterns to select objects or regions of interest within the field of view.^18, 46-47^ Alternatively, several investigators have explored measurements with line-excitation.^21, 48^ However, none of these previous patterned illumination studies incorporated intentional periodicity within the photobleach patterns that could subsequently integrate into FT-FRAP analysis. The closest work to the present study is arguably in early studies by Lanni and Ware, in which sinusoidal modulation of a bleach pattern was performed by passing the excitation beam through a grating.^49^ The apparatus was designed so that the image of the grating was at the focus of the sample, creating a photobleaching mask at high laser power. The subsequent fluorescence recovery was probed by translating the grating, producing a phase shift in the illumination pattern at the sample, which generated a periodic signal in the integrated fluorescence intensity over time as the grating was shifted. The integrated fluorescence signal was recorded on a single-channel detector and the Fourier components were analyzed to recover the diffusion coefficient. The conceptual foundation for the studies by Lanni and Ware is aligned with the principles undergirding the work presented herein. The difference lies in the tools that were used to realize the FT-FRAP measurement. Laser-scanning microscopy makes FT-FRAP a faster, higher SNR, measurement than similar techniques with different tools from decades ago.

### 4.4 FT-FRAP characterization of anomalous diffusion

Additional results were obtained to characterize anomalous diffusion in a viscous matrix. **Figure 5** shows FT-FRAP with a comb bleach pattern on FITC-polydextran (2 MDa) in 22 mg/mL hyaluronic acid (HA). HA is a glycosaminoglycan polymer that is found throughout the connective tissues of the body. The physiological functions in which HA is involved include lubrication, wound repair and cell migration. HA is known to increase viscosity when added to aqueous solution. In this experiment, the fluorescence recovery of the first three spatial harmonics of the comb bleach pattern was analyzed to characterize anomalous diffusion of FITC-polydextran in a solution with HA.

**Figure 5.**
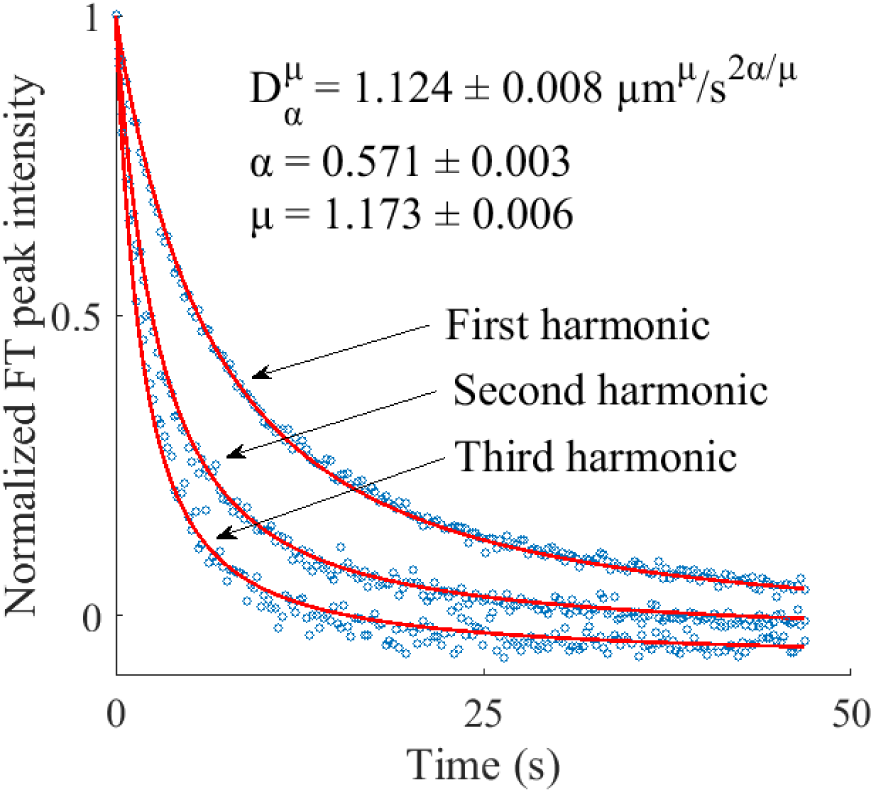
Harmonic analysis of anomalous diffusion with FITC-polydextran (2 MDa) in 22 mg/mL hyaluronic acid. Fluorescence recovery of the first, second, and third harmonics are fit to a modified Mittag-Leffler function to recover the anomalous diffusion coefficient, *D* = 1.124 ± 0.008 μm^*μ*^/s^2*α/μ*^, the subdiffusion parameter, *α* = 0.571 ± 0.003, and the Lévy flight parameter, *μ* = 1.173 ± 0.006. The recovered parameters reveal subdiffusive and Lévy flight behavior in the sample. The reported uncertainties are the standard deviations in the fit.

Several models for anomalous diffusion were considered. First, the three harmonics were fit to a model for systems with subdiffusive mobility, described in **Equation (16)**. Subdiffusion was considered because of possible binding and unbinding or association and dissociation of the FITC-polydextran with the HA matrix. However, analysis with this model did not produce a satisfactory fit because the data did not exhibit a quadratic dependence on spatial frequency. Second, the data were fit to a model for systems exhibiting Lévy flight, described in **Equation (18).** Lévy flight behavior was considered because of possible crowding or trapping by the HA acting as an obstacle to constrict free diffusion of the FITC-polydextran. While the Lévy flight model produced best-fit curves with a better match for the spatial frequency dependence, the shape of the best-fit curves was exponential and far from a good fit to the fluorescence recovery curves. Models incorporating just one of these two effects (subdiffusion and Lévy flight) were insufficient to describe the data.

Third, a global fit of the first three harmonics to a combined subdiffuion-Lévy flight model, described in **Equation (19)**, was performed to recover the anomalous diffusion coefficient *D* = 1.124 ± 0.008 μm^*μ*^/s^2*α*/*μ*^, the subdiffusive parameter *α* = 0.571 ± 0.003, and the Lévy flight parameter *μ* = 1.173 ± 0.006. The reported uncertainties are the standard deviations in the fit. The result of this fit is shown in **Figure 5**. The combined model was able to account for both the spatial frequency dependence of the harmonics and the shape of the recovery curves, which deviates from exponential. The value of *α* < 1 indicates that the sample exhibits a broad distribution in wait times (subdiffusion). Furthermore, the value for *μ* < 2 indicates that the sample exhibits a broad distribution in step lengths (Lévy flight).

The results of the analysis demonstrate that FT-FRAP can sensitively and precisely disentangle covariant parameters describing anomalous diffusion by simultaneously fitting to multiple harmonics acquired in parallel. The same analysis performed with only the first harmonic yields lower confidence in the recovered parameters: *D* = 1.9 ± 0.1 μm^*μ*^/s^2*α*/*μ*^, *α* = 0.69 ± 0.01, and *μ* = 1.57 ± 0.05. The reduction in precision likely arises from the increase in covariance in the single-harmonic fit; the recovered parameters are highly correlated (*D* & *α* = 0.976, *D* & *μ* = 0.995, and *α* & *μ* = 0.993, where ±1 corresponds to perfectly correlated / anti-correlated parameters). By comparison, the correlation coefficients obtained in the three-harmonic fit are much less significant: *D* & *α* = –0.204, *D* & *μ* = 0.286, and *α* & *μ* = 0.850. Comparison with a one-harmonic fit demonstrates that measuring diffusion at multiple length scales with FT-FRAP can substantially increase statistical confidence in the parameters recovered from fitting to an anomalous diffusion model by constraining the model to describe diffusion globally.

## 5. Conclusion

In this work, FT-FRAP with patterned illumination has been described theoretically and demonstrated experimentally to characterize normal and anomalous diffusion. Relative to conventional point-bleach FRAP, FT-FRAP has the advantages of mathematical simplicity, higher SNR, representative sampling, and multi-photon compatibility. Proof-of-concept measurements with a model system (FITC-polydextran) showed good agreement with theory. Flow was quantified using the phase of the real and imaginary components of the FT peaks in FT-FRAP. Anomalous diffusion was characterized by FT-FRAP through a global fit to multiple harmonics of the bleach pattern. Future fundamental work includes experiments to test the ability of FT-FRAP with patterned illumination to characterize heterogeneous samples, where diffusion varies across the field of view. FT-FRAP can also be implemented in a variety of application spaces including measurement of mobility in pharmaceutically relevant matrices, quantification of protein aggregation, and live-cell imaging.

## Author Contributions

AG and GS developed the theory; AG and GS designed the microscope; AG, CS, and MC built the microscope; AG, NT, and DH collected data; AG analyzed data; AG and GS wrote and edited the manuscript.

## Acknowledgements

The authors gratefully acknowledge funding for the present work from Eli Lilly & Company and from the National Science Foundation through an NSF-GOALI award (No. CHE-1710475).

## References

1. Loren, N.; Hagman, J.; Jonasson, J. K.; Deschout, H.; Bernin, D.; Cella-Zanacchi, F.; Diaspro, A.; McNally, J. G.; Ameloot, M.; Smisdom, N.; Nyden, M.; Hermansson, A. M.; Rudemo, M.; Braeckmans, K., Fluorescence recovery after photobleaching in material and life sciences: putting theory into practice. Q. Rev. Biophys. 2015, 48 (3), 323–387.

2. Reits, E. A. J.; Neefjes, J. J., From fixed to FRAP: measuring protein mobility and activity in living cells. Nat. Cell Biol. 2001, 3 (6), E145–E147.

3. Peters, R.; Peters, J.; Tews, K. H.; Bähr, W., A microfluorimetric study of translational diffusion in erythrocyte membranes. BBA - Biomembranes 1974, 367 (3), 282–294.

4. Vámosi, G.; Friedländer-Brock, E.; Ibrahim, S. M.; Brock, R.; Szöllősi, J.; Vereb, G., EGF Receptor Stalls upon Activation as Evidenced by Complementary Fluorescence Correlation Spectroscopy and Fluorescence Recovery after Photobleaching Measurements. Int. J. Mol. Sci. 2019, 20 (13), 3370.

5. Weiss, G. L.; Kieninger, A. K.; Maldener, I.; Forchhammer, K.; Pilhofer, M., Structure and Function of a Bacterial Gap Junction Analog. Cell 2019, 178 (2), 374-+.

6. Frojd, M. J.; Flardh, K., Apical assemblies of intermediate filament-like protein FilP are highly dynamic and affect polar growth determinant DivIVA in Streptomyces venezuelae. Mol. Microbiol. 2019, 112 (1), 47–61.

7. Verheyen, E.; van der Wal, S.; Deschout, H.; Braeckmans, K.; de Smedt, S.; Barendregt, A.; Hennink, W. E.; van Nostrum, C. F., Protein macromonomers containing reduction-sensitive linkers for covalent immobilization and glutathione triggered release from dextran hydrogels. J. Controlled Release 2011, 156 (3), 329–336.

8. Branco, M. C.; Pochan, D. J.; Wagner, N. J.; Schneider, J. P., Macromolecular diffusion and release from self-assembled beta-hairpin peptide hydrogels. Biomaterials 2009, 30 (7), 1339–1347.

9. Alvarez-Mancenido, F.; Braeckmans, K.; De Smedt, S. C.; Demeester, J.; Landin, M.; Martinez-Pacheco, R., Characterization of diffusion of macromolecules in konjac glucomannan solutions and gels by fluorescence recovery after photobleaching technique. International Journal of Pharmaceutics 2006, 316 (1-2), 37–46.

10. Kosto, K. B.; Deen, W. M., Diffusivities of macromolecules in composite hydrogels. Aiche J. 2004, 50 (11), 2648–2658.

11. Peeters, L.; Sanders, N. N.; Braeckmans, K.; Boussery, K.; Van de Voorde, J.; De Smedt, S. C.; Demeester, J., Vitreous: A barrier to nonviral ocular gene therapy. Invest Ophthalmol Visual Sci 2005, 46 (10), 3553–3561.

12. Remaut, K.; Sanders, N. N.; De Geest, B. G.; Braeckmans, K.; Demeester, J.; De Smedt, S. C., Nucleic acid delivery: Where material sciences and bio-sciences meet. Mater. Sci. Eng. R-Rep. 2007, 58 (3-5), 117–161.

13. Cherezov, V.; Liu, J.; Griffith, M.; Hanson, M. A.; Stevens, R. C., LCP-FRAP Assay for Pre-Screening Membrane Proteins for In Meso Crystallization. Crystal Growth & Design 2008, 8 (12), 4307–4315.

14. Joseph, J. S.; Liu, W.; Kunken, J.; Weiss, T. M.; Tsuruta, H.; Cherezov, V., Characterization of lipid matrices for membrane protein crystallization by high-throughput small angle X-ray scattering. Methods 2011, 55 (4), 342–349.

15. Xu, F.; Liu, W.; Hanson, M. A.; Stevens, R. C.; Cherezov, V., Development of an Automated High Throughput LCP-FRAP Assay to Guide Membrane Protein Crystallization in Lipid Mesophases. Crystal Growth & Design 2011, 11 (4), 1193–1201.

16. Chun, E.; Thompson, A. A.; Liu, W.; Roth, C. B.; Griffith, M. T.; Katritch, V.; Kunken, J.; Xu, F.; Cherezov, V.; Hanson, M. A.; Stevens, R. C., Fusion Partner Toolchest for the Stabilization and Crystallization of G Protein-Coupled Receptors. Structure 2012, 20 (6), 967–976.

17. Mazza, D.; Cella, F.; Vicidomini, G.; Krol, S.; Diaspro, A., Role of three-dimensional bleach distribution in confocal and two-photon fluorescence recovery after photobleaching experiments. Appl. Opt. 2007, 46 (30), 7401–7411.

18. Deschout, H.; Hagman, J.; Fransson, S.; Jonasson, J.; Rudemo, M.; Loren, N.; Braeckmans, K., Straightforward FRAP for quantitative diffusion measurements with a laser scanning microscope. Opt. Express 2010, 18 (22), 22886–22905.

19. Braeckmans, K.; Stubbe, B. G.; Remaut, K.; Demeester, J.; De Smedt, S. C., Anomalous photobleaching in fluorescence recovery after photobleaching measurements due to excitation saturation-a case study for fluorescein. JBO 2006, 11 (4), 13.

20. Braeckmans, K.; Peeters, L.; Sanders, N. N.; De Smedt, S. C.; Demeester, J., Three-dimensional fluorescence recovery after photobleaching with the confocal scanning laser microscope. Biophys. J. 2003, 85 (4), 2240–2252.

21. Braeckmans, K.; Remaut, K.; Vandenbroucke, R. E.; Lucas, B.; De Smedt, S. C.; Demeester, J., Line FRAP with the confocal laser scanning microscope for diffusion measurements in small regions of 3-D samples. Biophys. J. 2007, 92 (6), 2172–2183.

22. Davis, S. K.; Bardeen, C. J., Using two-photon standing waves and patterned photobleaching to measure diffusion from nanometers to microns in biological systems. Rev. Sci. Instrum. 2002, 73 (5), 2128–2135.

23. Smith, B. A.; McConnell, H. M., Determination of molecular-motion in membranes using periodic pattern photobleaching. Proc. Natl. Acad. Sci. U. S. A. 1978, 75 (6), 2759–2763.

24. Davoust, J.; Devaux, P. F.; Leger, L., Fringe pattern photobleaching, a new method for the measurement of transport-coefficients of biological macromolecules. EMBO J. 1982, 1 (10), 1233–1238.

25. Berk, D. A.; Yuan, F.; Leunig, M.; Jain, R. K., Fluorescence photobleaching with spatial Fourier-analysis - measurement of diffusion in light-scattering media. Biophys. J. 1993, 65 (6), 2428–2436.

26. Meyvis, T. K. L.; De Smedt, S. C.; Van Oostveldt, P.; Demeester, J., Fluorescence recovery after photobleaching: A versatile tool for mobility and interaction measurements in pharmaceutical research. Pharmaceutical Research 1999, 16 (8), 1153–1162.

27. Shi, C. C.; Kuo, J.; Bell, P. D.; Yao, H., Anisotropic Solute Diffusion Tensor in Porcine TMJ Discs Measured by FRAP with Spatial Fourier Analysis. Ann. Biomed. Eng. 2010, 38 (11), 3398–3408.

28. Tsay, T. T.; Jacobson, K. A., Spatial Fourier-analysis of video photobleaching measurements - principles and optimization. Biophys. J. 1991, 60 (2), 360–368.

29. Metzler, R.; Klafter, J., The random walk’s guide to anomalous diffusion: a fractional dynamics approach. Phys. Rep.-Rev. Sec. Phys. Lett. 2000, 339 (1), 1–77.

30. Klafter, J.; Blumen, A.; Shlesinger, M. F., Stochastic pathway to anomalous diffusion. Phys. Rev. A 1987, 35 (7), 3081–3085.

31. Wachsmuth, M.; Waldeck, W.; Langowski, J., Anomalous diffusion of fluorescent probes inside living cell nuclei investigated by spatially-resolved fluorescence correlation spectroscopy. J. Mol. Biol. 2000, 298 (4), 677–689.

32. Wong, I. Y.; Gardel, M. L.; Reichman, D. R.; Weeks, E. R.; Valentine, M. T.; Bausch, A. R.; Weitz, D. A., Anomalous diffusion probes microstructure dynamics of entangled F-actin networks. Phys. Rev. Lett. 2004, 92 (17), 4.

33. Banks, D. S.; Fradin, C., Anomalous diffusion of proteins due to molecular crowding. Biophys. J. 2005, 89 (5), 2960–2971.

34. Tolic-Norrelykke, I. M.; Munteanu, E. L.; Thon, G.; Oddershede, L.; Berg-Sorensen, K., Anomalous diffusion in living yeast cells. Phys. Rev. Lett. 2004, 93 (7), 4.

35. Daddysman, M. K.; Fecko, C. J., Revisiting Point FRAP to Quantitatively Characterize Anomalous Diffusion in Live Cells. J. Phys. Chem. B 2013, 117 (5), 1241–1251.

36. Heinze, K. G.; Costantino, S.; De Koninck, P.; Wiseman, P. W., Beyond photobleaching, laser illumination unbinds fluorescent proteins. The Journal of Physical Chemistry B 2009, 113 (15), 5225–5233.

37. Sokolov, I. M., Models of anomalous diffusion in crowded environments. Soft Matter 2012, 8 (35), 9043–9052.

38. Bouchaud, J.-P.; Georges, A., Anomalous diffusion in disordered media: statistical mechanisms, models and physical applications. Physics reports 1990, 195 (4-5), 127–293.

39. Sun, H.; Chen, W.; Chen, Y., Variable-order fractional differential operators in anomalous diffusion modeling. Physica A: Statistical Mechanics and its Applications 2009, 388 (21), 4586–4592.

40. Chen, W.; Sun, H.; Zhang, X.; Korošak, D., Anomalous diffusion modeling by fractal and fractional derivatives. Computers & Mathematics with Applications 2010, 59 (5), 1754–1758.

41. Saxton, M. J., Anomalous diffusion due to obstacles - a Monte-Carlo study. Biophys. J. 1994, 66 (2), 394–401.

42. Bracewell, R. N.; Bracewell, R. N., The Fourier transform and its applications. McGraw-Hill New York: 1986; Vol. 31999.

43. Muir, R. D.; Sullivan, S. Z.; Oglesbee, R. A.; Simpson, G. J., Synchronous digitization for high dynamic range lock-in amplification in beam-scanning microscopy. Rev. Sci. Instrum. 2014, 85 (3), 033703.

44. Garrappa, R., Numerical evaluation of two and three parameter Mittag-Leffler functions. SIAM J. Numer. Anal. 2015, 53 (3), 1350–1369.

45. vanderSchaaf, A.; vanHateren, J. H., Modelling the power spectra of natural images: Statistics and information. Vision Res. 1996, 36 (17), 2759–2770.

46. Wedekind, P.; Kubitscheck, U.; Peters, R., Scanning microphotolysis - a new photobleaching technique based on fast intensity modulation of a scanned laser-beam and confocal imaging. Journal of Microscopy-Oxford 1994, 176, 23–33.

47. Blumenthal, D.; Goldstien, L.; Edidin, M.; Gheber, L. A., Universal Approach to FRAP Analysis of Arbitrary Bleaching Patterns. Sci. Rep. 2015, 5.

48. Wedekind, P.; Kubitscheck, U.; Heinrich, O.; Peters, R., Line-scanning microphotolysis for diffraction-limited measurements of lateral diffusion. Biophys. J. 1996, 71 (3), 1621–1632.

49. Lanni, F.; Ware, B. R., Modulation detection of fluorescence photobleaching recovery. Rev. Sci. Instrum. 1982, 53 (6), 905–908.

